# A biophysical minimal model to investigate age-related changes in CA1 pyramidal cell electrical activity

**DOI:** 10.1101/2022.07.01.498486

**Authors:** Erin C. McKiernan, Marco A. Herrera-Valdez, Diano F. Marrone

## Abstract

Aging is a physiological process that is still poorly understood, especially with respect to effects on the brain. There are open questions about aging that are difficult to answer with an experimental approach. Underlying challenges include the difficulty of recording *in vivo* single cell and network activity simultaneously with submillisecond resolution, and brain compensatory mechanisms triggered by genetic, pharmacologic, or behavioral manipulations. Mathematical modeling can help address some of these questions by allowing us to fix parameters that cannot be controlled experimentally and investigate neural activity under different conditions. We present a biophysical minimal model of CA1 pyramidal cells (PCs) based on general expressions for transmembrane ion transport derived from basic thermodynamical principles. The model allows directly varying the contribution of ion channels by changing their number. By analyzing the dynamics of the model, we find parameter ranges that reproduce the variability in electrical activity seen in PCs. In addition, increasing the L-type Ca^2+^ channel expression in the model reproduces age-related changes in excitability that are qualitatively and quantitatively similar to those observed in PCs from aged animals.

## 1 Introduction

As we age, our brains undergo many changes [1, 2], but we understand relatively little about these and their effects on neural function. What does normal neurophysiological aging look like and what are the various stages? How does the electrical activity of neurons change and what are the biophysics underlying those changes? How do aging neurons respond to input from other cells? Answering these questions is not just fundamental to understanding aging as a neurophysiological process, but also to understanding how this process may be altered in age-related disorders of clinical importance such as Alzheimer’s [3] and Parkinson’s [4] disease.

Many aging studies have focused on the hippocampus, an area of the brain involved in learning, memory formation, and spatial processing [1, 2]. Aged rats [5–7] and humans [8] show impaired learning of hippocampal-dependent spatial tasks. Long-term potentiation (LTP), a proposed physiological substrate of memory formation, has been investigated in the hippocampus and its induction and maintenance shown to be impaired in aged rats [9, 10]. A short-term form of plasticity, frequency potentiation/facilitation (FP/FF), is also impaired in hippocampal pyramidal cells (PCs) from aged rats and correlates with learning deficits [11].

Plasticity changes and behavioral impairments may result in part from altered Ca^2+^ signaling in aged neurons [1, 2]. Compared to CA1 PCs from young animals, PCs from aged animals show larger and longer post-burst afterhyperpolarizations (AHPs) [12–14]. AHPs are mediated by Ca^2+^-dependent K^+^ currents, which can act like brakes on the electrical activity of CA1 PCs [15, 16]. As a result, PCs show increased spike frequency adaptation and fire fewer action potentials (APs) in response to acute stimuli or during bursting activity [17–19]. Larger AHPs are associated with increased intracellular Ca^2+^, mediated in part by Ca^2+^ entry via L-type channels [14, 18, 20, 21]. Aged animals show increases in L-type channel expression and/or channel density at the plasma membrane [22–25]. Animals with higher Ca^2+^ channel density perform poorly in spatial tasks [23], while blockers of L-type channels can restore learning and plasticity in older animals [26, 27].

It is not well understood how changes in ion channel gene expression and hippocampal PC excitability may affect neuron responsiveness and microcircuit output. In part, this is due to challenges inherent in performing the needed experiments. Single PCs are difficult to access in intact animals where hippocampal microcircuit function is preserved. It is also difficult to tease apart the influence of the many different neurophysiological factors that change during aging. Mathematical modeling provides means to understand more about the effects of aging on hippocampal cellular excitability by controlling factors we cannot control experimentally.

Our previous work shows that mathematical expressions for different passive and active ion transport mechanisms can be derived from first principles of thermodynamics [28, 29] using a common functional form [30]. This results in a realistic representation of ionic flow across the membrane, and allows the model to reproduce phenomena such as rectification of ion currents seen in recordings. We present a model that reproduces the diversity of firing patterns in CA1 PC recordings, including adaptive firing, stimulus-induced bursting, and spontaneous bursting [31]. In addition, we reproduce several electrophysiological characteristics of aging by varying the expression of Ca^2+^ channels in the model, and make predictions about bursting activity in aged CA1 PCs, which to our knowledge has not been reported. We believe this model is ideal to further study the effects of various biophysical changes in CA1 PCs during aging, as well as potentially forming the basis for biophysical, yet computationally inexpensive, network models.

## 2 Methods

### 2.1 Model

To simulate the electrical activity of CA1 PCs, we used an extended version of a two-dimensional, biophysical model previously developed and characterized by two of the present authors [28–30]. The equations for the ionic currents are derived from first principles of thermodynamics. Previous modeling studies have shown that to reproduce firing behaviors such as spike frequency adaptation and bursting, the minimum number of variables is three [32, 33]. In particular, Ca^2+^ dynamics are important for producing adaptation and burst firing in CA1 PCs (for review see [31]). The model dynamics are therefore described by three ordinary differential equations for the time-dependent changes in the transmembrane potential (*v*, in mV), the proportion of open K^+^ channels (*w* in [0,1]), and the intracellular Ca^2+^ concentration (*c*, in *μ*M), respectively [30]. Based on a well-known relationship between voltage-dependent activation of delayed rectifier K^+^ channels and inactivation of Na^+^ channels, *w* also represents the proportion of inactivated Na^+^ channels [34, 35].

It is assumed that the membrane potential changes due to currents produced by ions transported across the membrane. We take into account currents mediated by a voltage-gated, inactivating Na^+^ channel (*I*_*NaT*_), voltage- and Ca^2+^-gated K^+^channels (*I*_*DK*_ and *I*_*SK*_, respectively), and a Na^+^/K^+^-ATPase (*I*_*NaK*_). The time-dependent change in membrane potential can be written as

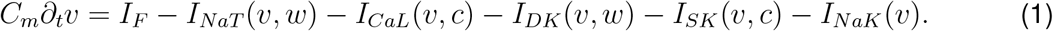

with *∂*_*t*_ representing the instantaneous change with respect to time. *C*_*m*_ (pF) is a constant representing the change in the density of charge around the membrane with respect to voltage, typically referred to as membrane capacitance in models based on electrical circuits [36]. Based on recordings from rat CA1 PCs, it is assumed that *C*_*m*_ = 25 pF [37].

All the currents in Eq. (1) are modeled using the same generic functional form, a product

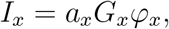

*x* ∈ {*NaT, CaL, DK, SK, NaK*}, where *a*_*x*_ is a whole-cell current amplitude (pA), *G*_*x*_ is a gating term (between 0 and 1), and *φ*_*x*_ is an adimensional term describing the driving force for the transmembrane flux (Table 1). The terms *a*_*x*_ = *s*_*x*_*N*_*x*_, *x* ∈ {*NaT, CaL, DK, SK, NaK*} are whole-cell current amplitudes with *s*_*x*_ (pA) representing the current flowing through a single channel (or pump), and *N*_*x*_ representing the number of membrane proteins mediating the current (e.g. number of K^+^ channels). Of interest, *s*_*x*_ is ∼1 pA for most voltage-gated channels [38], and is ∼5-10 pA for SK channels [39].

**Table 1:** Transport mechanisms included in the model.

| Transport mechanism | Current | Amplitude ($a$) | Gating ($G$) | Flux $\varphi$ |
| --- | --- | --- | --- | --- |
| Transient $Na^+$ channels | $I_{NaT}(v, w)$ | $a_{Na}$ | $S_m(v)(1 - w)$ | $\varphi_{Na}(v)$ |
| L-type $Ca^{2+}$ channels | $I_{CaL}(v, c)$ | $a_{Ca}$ | $S_n(v)$ | $\varphi_{Ca}(v)$ |
| Delayed rectifier $K^+$ channels | $I_{DK}(v, w)$ | $a_{DK}$ | $w$ | $\varphi_K(v)$ |
| SK $Ca^{2+}$-dependent $K^+$ channels | $I_{SK}(v, c)$ | $a_{SK}$ | $H_{SK}(c)$ | $\varphi_K(v)$ |
| $Na^+/K^+$ pumps | $I_{NaK}(v)$ | $a_{NaK}$ | 1 | $\varphi_{NaK}(v)$ |

All ion fluxes are given by a product of the form $I_x = a_x G_x \varphi_x$ , where $a_x$ , $G_x$ , and $\varphi_x$ represent, respectively, the amplitude (normalized by membrane capacitance), gating, and driving force terms for the flux. $G_{NaK} = 1$ represents saturation of the $Na^+/K^+$ pumps. Note that inactivation of $Na^+$ channels is also represented by $w$ [34, 35], so the proportion of non-inactivated $Na^+$ channels is $1 - w$ .

Assuming that none of the currents in the model exhibit rectification [30], which agrees with recordings of the included currents, the adimensional component of the transmembrane flux can be simplified and written as

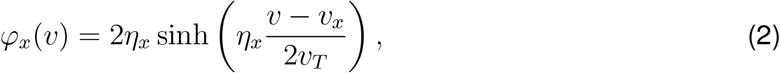

where *η*_*x*_ represents the number of charges transported across the membrane in a single transport event, and *v*_*x*_ is the reversal potential for the current, *x* ∈ {*NaT, CaL, DK, SK, NaK*}. For channels, *v*_*x*_ is the Nernst potential for the ion [30]. Of note, *η*_*x*_ = 1 for *x* ∈ {*DK, SK, NaK*} and *η*_*NaT*_ = −1 [30], which gives

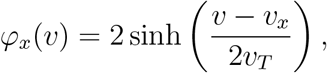

for *x* ∈ {*NaT, DK, SK, NaK*}. For Ca^2+^ channels, the total charge transported by one ion crossing the membrane from the extracelular space is *η*_*CaL*_ = −2, so

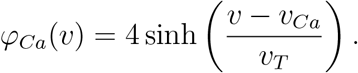

The driving force for flux is assumed to be the same for DK and SK channels. Therefore, the label *K* is used for both fluxes from here on. The thermal potential *v*_*T*_ = *kT/q* (mV), where *k* is Boltzmann’s constant (mJ/^*o*^K), *T* is the absolute temperature (^*o*^K), and *q* is the elementary charge (Coulombs). The Boltzmann constant can be thought of as a scaling factor between macroscopic (thermodynamic temperature) and microscopic (thermal energy) physics [40]. The reversal potentials for the different currents depend on the Nernst potentials for each ion, as given by

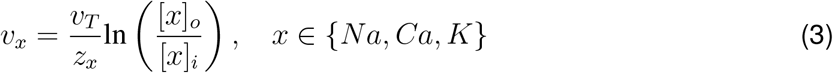

where *z*_*x*_ is the ion valence and [*x*]_*o*_ and [*x*]_*i*_ are the ion concentrations outside and inside the cell, respectively. The reversal potential for the Na^+^/K^+^-ATPase is given by *v*_*NaK*_ = *v*_*AT P*_ + 3*v*_*Na*_ − 2*v*_*K*_ [30]. The Nernst potentials for Na^+^ and K^+^ are assumed to be constant, but *v*_*Ca*_ varies because the intracellular Ca^2+^ concentration is a state variable in the model.

#### Gating

The auxiliary functions describing voltage-dependent activation are given by

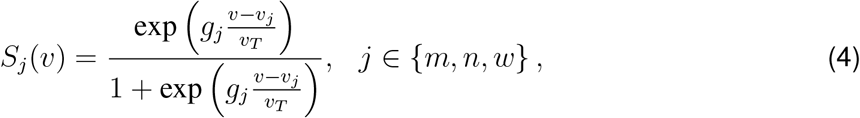

where *g*_*j*_ controls the steepness of the activation curve for Na^+^ (*m*), Ca^2+^ (*n*), or DK (*w*) channels, and *v*_*j*_ represents the half-activation voltage for those channels. The function

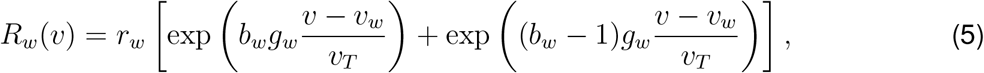

describes the voltage-dependence of the rate of activation of the DK channels. The parameters *r*_*w*_ and *b*_*w*_ represent the recovery rate and the asymmetry in the gating relative to voltage that biases the time constant for the gating process, respectively.

The dynamics for the proportion of activated DK channels, *w*, are assumed to be logistic,

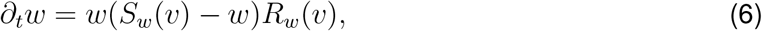

which yields better fits and is more consistent with the dynamics of activation in channel populations recorded in voltage-clamp experiments (e.g., see the activation curves in [41–43]).

The gating of the SK channel is not voltage-dependent, but instead depends on intracellular Ca^2+^ binding. Its activation is modeled using a Hill equation that depends on the intracellular concentration of Ca^2+^, as used to fit data from channel recordings [44]:

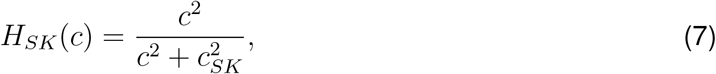

where *c*_*SK*_ represents the half-activation Ca^2+^ concentration for the SK channels, with a reported value of 0.74 *μ*M (740 nM) [39, 44].

For the dynamics of intracellular Ca^2+^, we assume recovery toward a steady state *c*_∞_ at a rate *r*_*c*_, with increments caused by the Ca^2+^ current *I*_*Ca*_ [35],

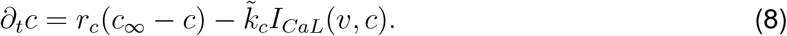

The term 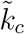 in equation (8) is a conversion factor (*μ*M/pCoul) that accounts for the effect of Ca^2+^ flux across the membrane on the intracellular Ca^2+^ concentration.

The term *I*_*F*_ represents a stimulus *forcing* the membrane; that is, current from an electrode, or time-dependent fluctuations from the local field potential. LFP activity is simulated by replacing the term *I*_*F*_ with a time-dependent, Ornstein-Uhlenbeck (OU) process with amplitude *a*_*F*_ (*t*) (pA). The mean is represented by *μ*_*F*_ (pA) (drift term) [45] given by [46]

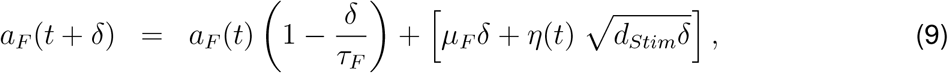

where *δ* is a small time step, *τ*_*F*_ is a relaxation time, and *η*(*t*) is an independent white noise process with zero-mean and unit standard deviation. The mean is set close to the rheobase for the model PCs (∼50 pA). The process has a variance 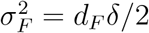 (pA), which means *d*_*F*_ can be approximated if an estimation of the variance of the current *a*_*F*_ is available [47, 48].

#### Change of variables to obtain numerical solutions

To simplify the numerics, we change variables

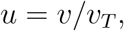

and adjust all voltages accordingly as

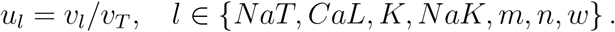

The new equation for the normalized voltage is

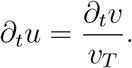

To simplify the notation and reduce the number of operations during the numerical integration, we also reparametrize the amplitudes as

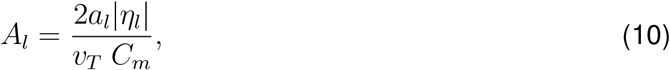

for *l* ∈ {*NaT, CaL, K, NaK*}, in units of 1/ms. Similarly, the activation functions for the Na^+^, Ca^2+^, and DK currents can also be rewritten respectively as

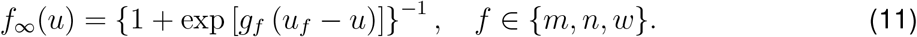

The result is a new equation of the form

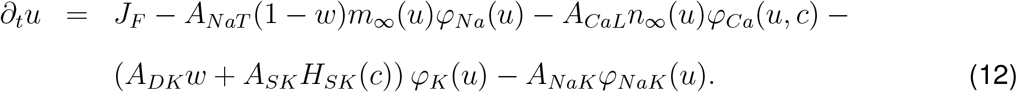

The term *J*_*F*_ (1/ms) is the input current *I*_*F*_ (pA) divided by *v*_*T*_ *C*_*m*_. After the change in variables and the normalization of the current amplitudes, equation (8) changes to

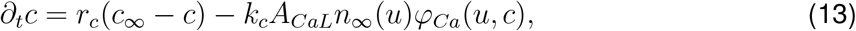

where 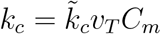.

### 2.2 Parameters

The currents were modeled to fit as closely as possible the biophysical properties of those carried by channel variants expressed in mammalian neurons, and specifically CA1 PCs, where data are available. The DK current is based on that mediated by K_*v*_2.1 channels, the predominant channel underlying the delayed rectifier current in rat hippocampal neurons [49]. The L-type Ca^2+^ current is based on that carried by Ca_*v*_1.2 (class C) channels, the predominant L-type channel isoform expressed in rat brain [50]. Additional details about the parameters can be found in Table 2.

**Table 2:** Constants and parameters.

| parameter | description | value | units | reference |
| --- | --- | --- | --- | --- |
| $k$ | Boltzmann's constant | $1.381\text{e}^{-20}$ | mJ/K | physical constant [38] |
| $q$ | elementary charge | $1.602\text{e}^{-19}$ | C | physical constant [38] |
| $T$ | absolute temperature | 273.15<br>+ 37 | K | adjusted to mammalian body temperature of 37°C [38] |
| $a_{NaT}$ | amplitude of transient $\text{Na}^+$ current | 1000-2300 | pA | set to produce currents of $\sim 2\text{-}7$ nA, in range recorded in CA1 PCs from rats [53] and guinea pigs [54] |
| $a_{CaL}$ | amplitude of L-type $\text{Ca}^{2+}$ current | 25 or 50 | pA | set to produce currents of $\sim 2\text{-}3$ nA or $\sim 5\text{-}6$ nA as recorded in young and aged CA1 PCs, respectively [20] |
| $a_{DK}$ | amplitude of delayed rectifier $\text{K}^+$ current | 6000-8000 | pA | set to produce currents of $\sim 6\text{-}9$ nA, in range recorded from HEK cells expressing rat Kv2.1 and $I_K$ in hippocampal neurons [55] |
| $a_{SK}$ | amplitude of $\text{Ca}^{2+}$-dependent $\text{K}^+$ current | 300-1600 | pA | set to produce currents of $\sim 100\text{-}800$ pA, depending on $\text{Ca}^{2+}$ concentration, as recorded in SK-transfected cells [56] |
| $a_{NaK}$ | amplitude of $\text{Na}^+/\text{K}^+$-ATPase current | 10-23 | pA | set to produce currents of $\sim 70\text{-}190$ pA, similar to but on high end of range recorded in hippocampal PCs [57] |
| $\eta_x$ | charge moved across the membrane in a single transport event<br>$x \in \{NaK, DK, SK\}$ | 1 | – | 1 net positive charge moving outward [30] |
| $\eta_{NaT}$ | charge transported across the membrane by $NaT$ channels | -1 | – | 1 net positive charge moving inward [30] |
| $\eta_{CaL}$ | charge transported across the membrane by Ca channels | -2 | – | 2 net positive charges moving inward [30] |
| $v_{Na}$ | Nernst potential for $\text{Na}^+$ | 60 | mV | in range reported for mammalian cells [58] |
| $[Ca]_o$ | extracellular $\text{Ca}^{2+}$ concentration (intracellular varies) | 1.5 | mM | in range reported for mammalian neural tissue [59] |
| $v_{Ca}$ | Nernst potential for $\text{Ca}^{2+}$ | variable; baseline $\sim 128$ | mV | in range reported for mammalian cells [58]; varies since intracellular $\text{Ca}^{2+}$ concentration is a model variable |
| $v_K$ | Nernst potential for $\text{K}^+$ | -89 | mV | in range reported for mammalian cells [58] |
| $v_{ATP}$ | Nernst potential for ATP | -420 | mV | value used in model of mammalian heart cells and based on fit to data [60] |
| $v_{NaK}$ | Nernst potential for $\text{Na}^+/\text{K}^+$-ATPase | -62 | mV | calculated based on the Nernst potentials for ATP, $\text{Na}^+$, and $\text{K}^+$, and a 3:2 stoichiometry, respectively [61]; $v_{NaK} = 3v_{Na} - 2v_K - v_{ATP}$ |
| $r_w$ | rate of activation of delayed rectifier $K^+$ current | 1.0-1.8 | ms | fit so that the duration of the action potential is approximately 2 ms [62] |
| $s_w$ | asymmetry of time constant of delayed rectifier $K^+$ current | 0.3 | - | based on fit; if higher (0.5-0.7) APs are the wrong shape and do not ride on sufficient plateau potential compared to recordings |
| $v_m$ | half-activation potential of $Na^+$ current | -19 | mV | in range reported for transient $Na^+$ channels in CA1 PCs [63, 64] |
| $v_n$ | half-activation potential of $Ca^{2+}$ current | 3 | mV | in range recorded for high-voltage activated $Ca^{2+}$ currents in rat CA1 PCs [65]; see also recordings from oocytes [66] or HEK cells [67] expressing $Ca_v1.2$ channels |
| $v_w$ | half-activation potential of delayed rectifier $K^+$ current | -1 | mV | in range reported for rat Kv2.1 channels expressed in COS-1 cells [49] |
| $c_{SK}$ | half-activation $Ca^{2+}$ concentration for SK current | $7.4e^{-4}$ | mM | based on recordings from oocytes expressing rat SK channel variant [44] |
| $g_m$ | activation slope of $Na^+$ current | 5.0 | - | in range reported for $Na^+$ current in mouse [68] and rat [69] CA1 neurons |
| $g_n$ | activation slope of $Ca^{2+}$ current | 5.0 | - | in range reported for $Ca_v1.2$ expressed in HEK cells [70] |
| $g_w$ | activation slope of delayed rectifier $K^+$ current | 3.8 | - | fit to data from rat brain delayed rectifier channels [71] |
| $c_\infty$ | minimum intracellular $Ca^{2+}$ concentration | $1e^{-4}$ | mM | equivalent to 100 nM, approximate resting intracellular $Ca^{2+}$ concentration in rat CA1 PCs [17, 72, 73] |
| $r_c$ | intracellular $Ca^{2+}$ removal rate constant | $1e^{-3}$ to $5e^{-3}$ | $ms^{-1}$ | adjusted to produce $Ca^{2+}$ dynamics as recorded in rat CA1 PCs [72] |
| $k_c$ | conversion factor to calculate effect of $Ca^{2+}$ current on intracellular $Ca^{2+}$ concentration | $3e^{-6}$ to $6e^{-6}$ | mM | adjusted to produce $Ca^{2+}$ dynamics as recorded in rat CA1 PCs [72] |

Wherever possible, model parameters were taken from studies in rodent (mice and rat) hippocampal CA1 PCs. If data were not available, we obtained parameters from other types of mammalian cell, or from studies of mammalian ion channels in expression systems like *Xenopus* oocyte. Physical constants and other parameters we would not expect to vary, such as the intra- and extracellular concentrations of ions or the cellular capacitance, were fixed. Biophysical properties of the ion channels, such as their half-activation voltages, were also fixed. The parameters we varied were primarily those corresponding to maximum current amplitudes, which can change acutely due to modulation or channel phosphorylation [51, 52], or chronically due to changes in ion channel expression that occur with age [22, 24].

By exploring the model through parameter variations, we were able to find parameter sets that produced different firing patterns, such as adaptive firing, conditional bursting, and spontaneous bursting. These firing patterns are described in more detail in the Results section, but the respective parameter sets are included in Table 3 for ease of comparison.

**Table 3:**
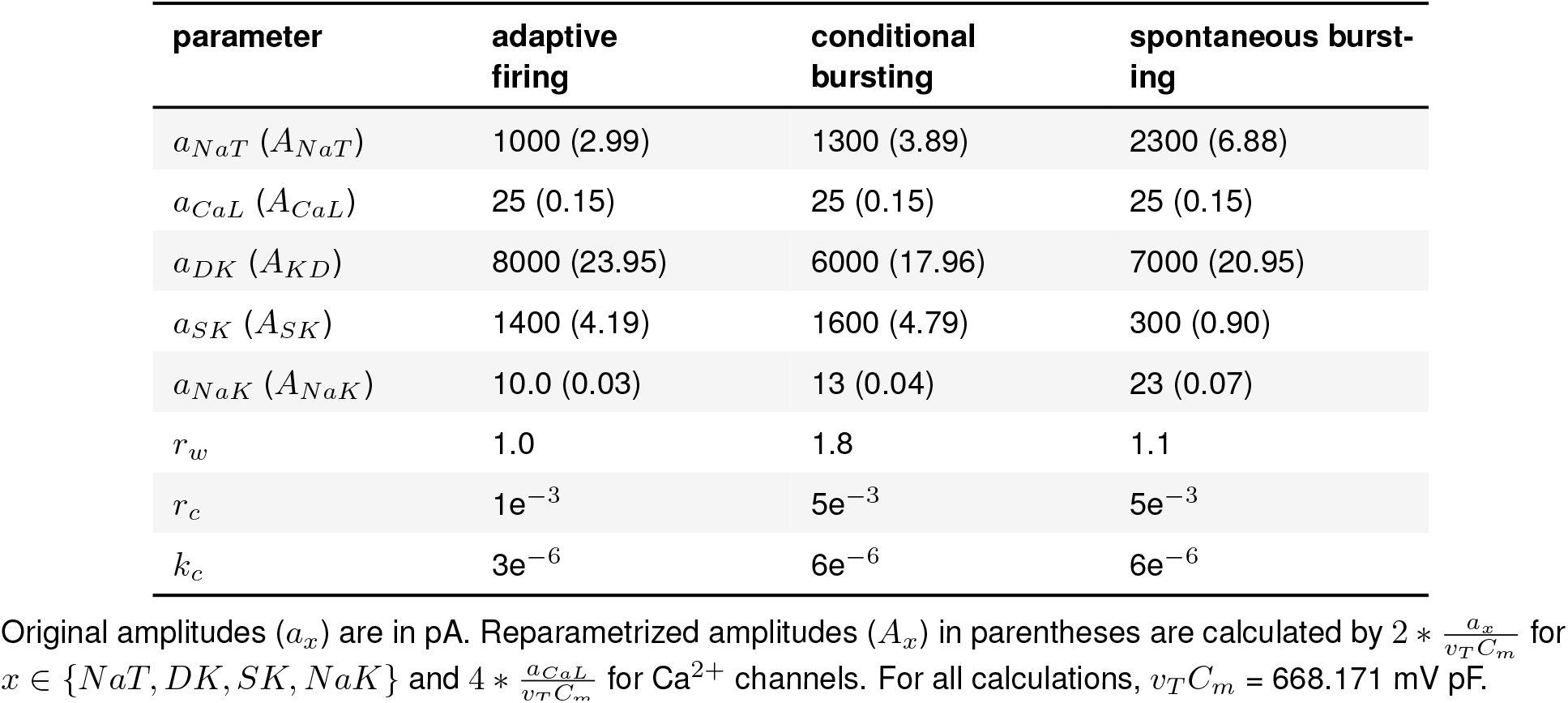
Parameters used to produce different firing patterns in the young PC model. See equation (10).

### 2.3 Simulations

All code was written in Python 3.7.4 and run on MacBook Pro laptops with 2.9 GHz Intel Core i5 processors. Simulations were performed using functions from the Python library NumPy [74]. Figures were produced with the Python library Matplotlib [75]. OU processes were simulated using the pyprocess (pypi.org/project/pyprocess) module [76].

### 2.4 Resource availability

Resources generated by this study (code, figures, and manuscript files) are available via GitHub (github.com/emckiernan/agingCA1) and archived via Zenodo (doi.org/10.5281/zenodo.6788229). To facilitate reuse, resources are shared under open licenses (see the license file in our GitHub repository). To promote reproducibility, Python code is embedded in a Jupyter notebook [77] that explains the code, how to use it, and how to generate the figures herein, plus additional ones.

### 2.5 Study design

While aged cells display a number of biophysical changes, we focused on their Ca^2+^ channel expression. Aged CA1 PCs show an increase in the number of functional transmembrane L-type Ca^2+^ channels [22–24]. In particular, CA1 PCs from aged rats have increased expression of Ca_*v*_1.2 at the plasma membrane [25]. With these results in mind, we decided to simulate one aspect of aging by changing the number of Ca_*v*_1.2 channels in our model membrane. We asked the question, is a change in Ca_*v*_1.2 expression sufficient to reproduce the various changes in excitability, such as increased spike frequency adaptation, observed experimentally in aged CA1 PCs? In addition, CA1 PCs are known to burst [31], but we are not aware of studies comparing the bursting patterns in young versus aged animals. Therefore, we used our model to also investigate the effects of altered Ca_*v*_1.2 channel expression on bursting activity. In all the following simulations, once the models have been tuned to a specific firing pattern, young and aged model PCs (hereafter referred to as yPC and aPC, respectively) are identical with respect to every parameter except the maximum amplitude of their L-type Ca^2+^ current, which is set to produce currents of ∼2-3 nA or ∼5-6 nA to match the magnitude of currents seen in recordings of CA1 PCs from young and aged animals, respectively [20].

## 3 Results

CA1 PCs display diverse firing patterns, ranging from repetitive adaptive spiking to stimulus-induced or spontaneous bursting (for review see [31]). Thus, to represent these cells, our model must reproduce this diversity, as well as age-related effects on firing reported in the literature.

### 3.1 Modeling age-related changes in spike frequency adaptation

Many CA1 PCs respond to square-pulse current injection by firing several early spikes followed by adaptation which slows the frequency of firing [17, 18, 78]. To generate this firing pattern, we set the ionic currents to be the same amplitude range as observed in recordings of young adult CA1 PCs, with amplitudes for the Na^+^ and Ca^2+^ currents at ∼2-3 nA, the DK current approximately double the inward cationic currents, and the SK current at ∼400-700 pA, depending on the intracellular Ca^2+^ concentration (see Tables 2 and 3). This balance of ionic currents successfully generates adaptive firing similar to recordings (Fig. 1, top panel; and additional figure in our Jupyter notebook).

**Figure 1:**
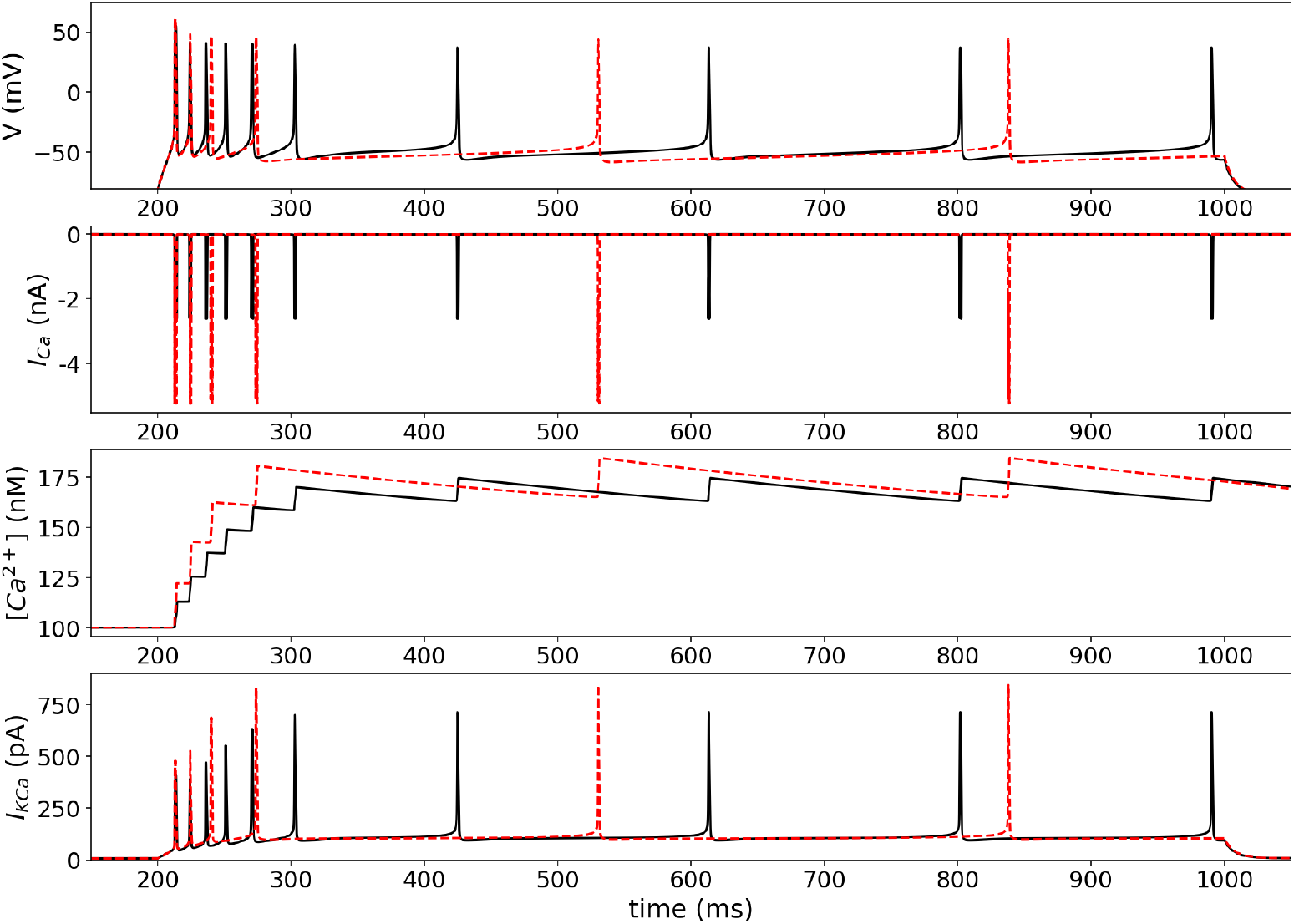
Adaptive firing in the yPC (solid black traces) versus aPC (dashed red traces) in response to a 800 ms square-pulse stimulation (top panel). Corresponding Ca^2+^ currents, intracellular Ca^2+^ concentration, and SK currents are in the 2nd, 3rd, and 4th panels, respectively.

Studies show that adaptation is more pronounced in aged than young animal cells, leading to a shorter initial period of fast spiking, followed by fewer spikes or complete cessation of firing [17–19, 79]. To compare the young (yPC) and aged (aPC) model cells, all parameters were fixed except for the maximum amplitude of the L-type Ca^2+^ current, which was set to produce currents of ∼2-3 nA (young) or ∼5-6 nA (aged), based on recordings [20]. This difference in the Ca^2+^ current causes the rate of firing in the first ∼100 ms (early firing) to decrease from 60 Hz in the yPC to 40 Hz in the aPC (Fig. 1). Firing for the remaining stimulation time (late firing) is also affected, decreasing from ∼6 Hz to ∼3 Hz in the yPC versus aPC, respectively. The effect and frequencies are similar to those seen in recordings of CA1 PCs in young and old rabbits [18].

Examining the Ca^2+^ and SK dynamics during the response reveals the mechanisms underlying the stronger adaptation in the aPC (Fig. 1, 2nd through 4th panels). The first two spikes occur nearly simultaneously in the two model cells. However, the larger increase in intracellular Ca^2+^ in the aPC induces a larger SK current, which in turn slows the cell’s firing. The aPC falls behind the yPC by the third spike, and then slows its firing further as the response continues.

**Figure 2:**
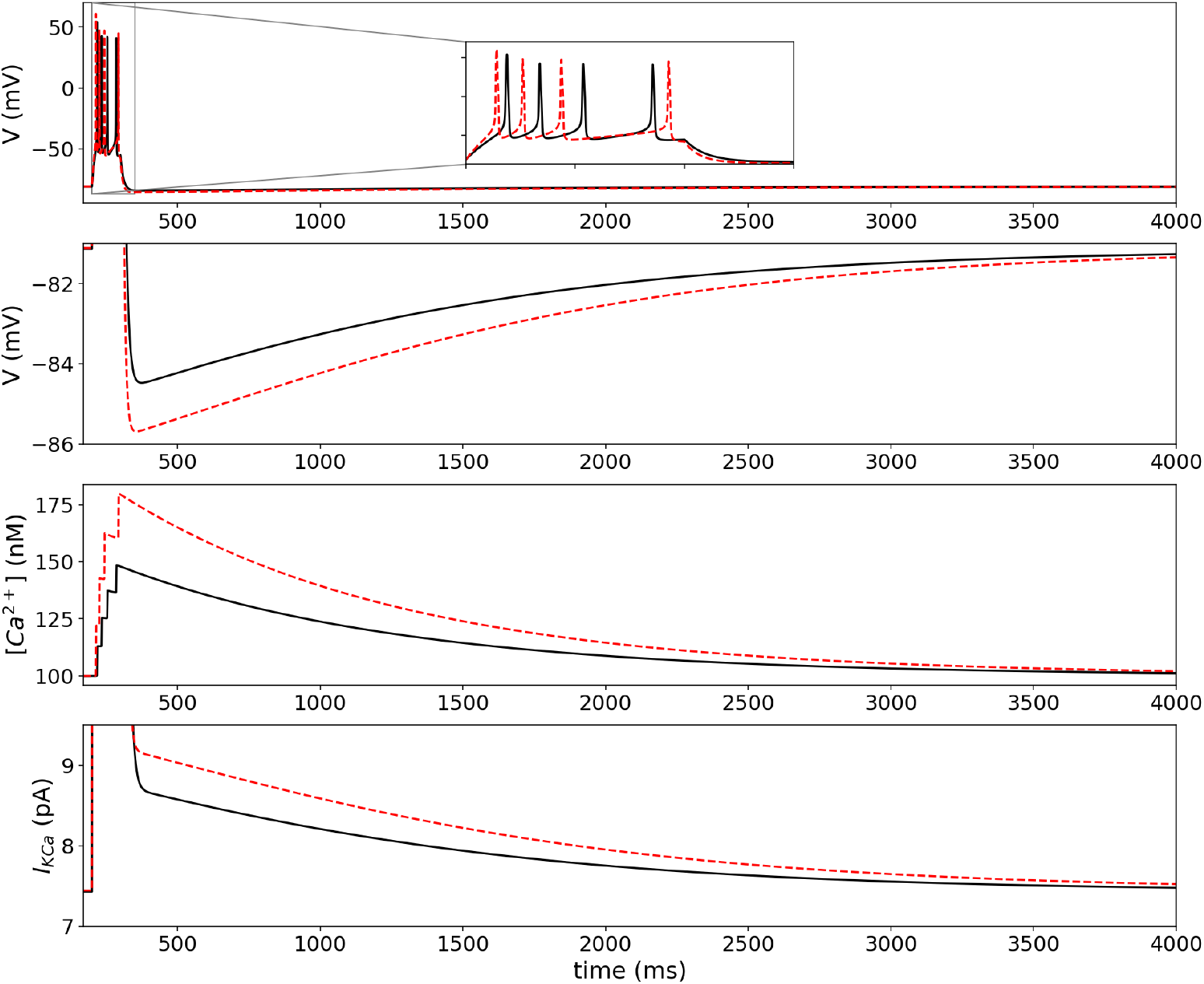
Responses of the yPC (solid black traces) and aPC (dashed red traces) to 100 ms pulse (top panel). Current amplitude adjusted to produce 4 spikes in each cell (yPC: 112 pA; aPC: 142 pA). Voltage zoom in 2nd panel shows AHPs in detail. Corresponding Ca^2+^ concentrations and SK currents are in the 3rd and 4th panels, respectively.

### 3.2 Modeling age-related changes in AHPs

AHP generation has been studied in CA1 PCs [80], particularly in the context of aging [17, 18, 79, 81]. To induce AHPs, we kept the same parameters as in the previous simulations. We then stimulated model PCs with a 100 ms square pulse of sufficient amplitude to generate a burst of 4 APs (Fig. 2). The AHPs produced under these conditions in the yPC have a peak amplitude of 3-4 mV, similar to recordings [14, 81]. The aPC required 30 pA more current than the yPC to fire the same number of spikes. However, the aPC fires earlier than the yPC due to its increased Ca^2+^ current (Fig. 2, top panel inset). The aPC generates an AHP 1-2 mV larger than seen in the yPC (Fig. 2, 2nd panel), similar to the difference observed in recordings between young and aged cells [14, 17, 18]. In the model, this larger AHP is due to an increased accumulation of Ca^2+^ in the aPC, which in turn produces a larger SK current (Fig. 2, 3rd and 4th panels).

**Figure 3:**
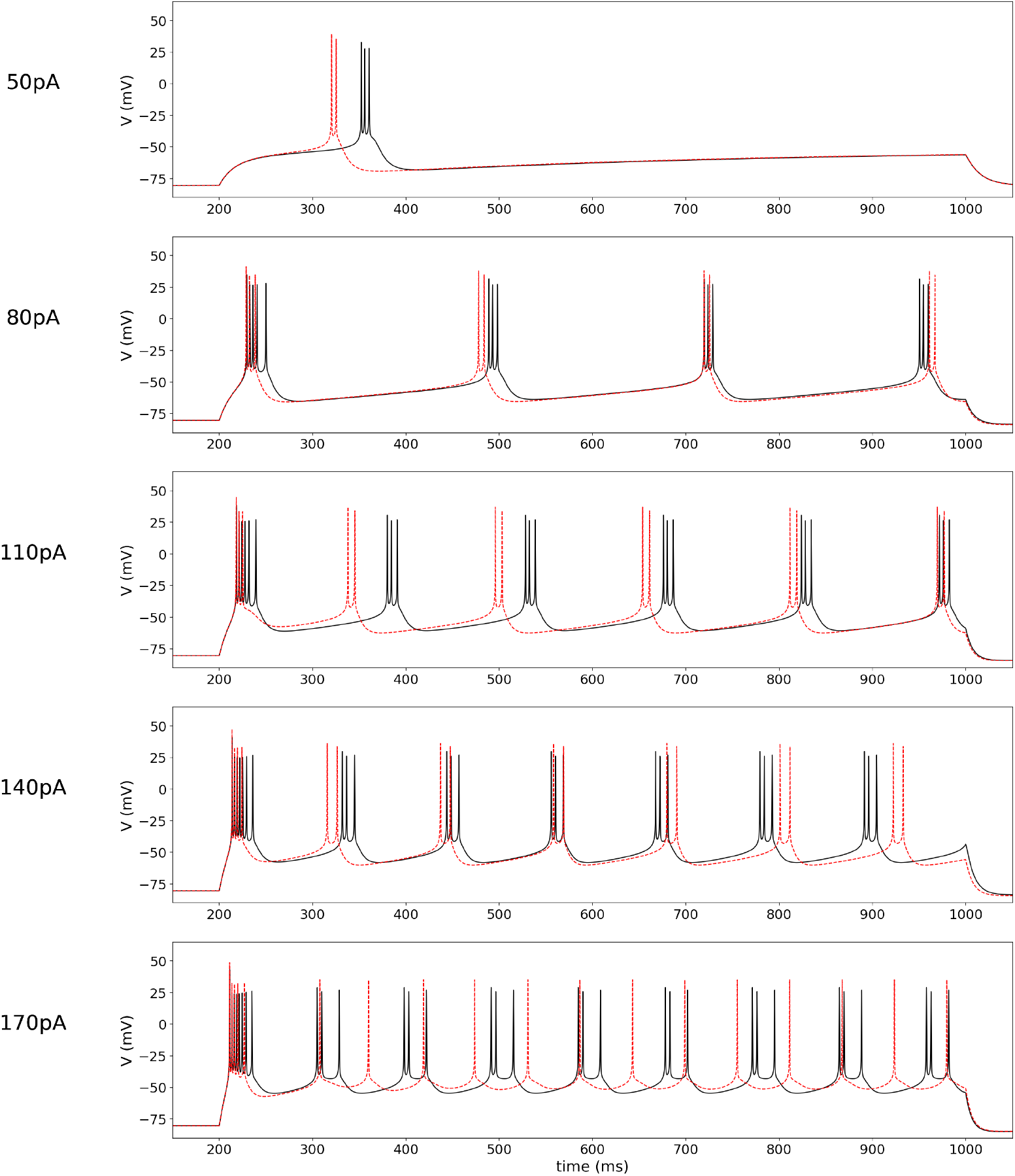
Bursting in the yPC (solid black traces) and aPC (dashed red traces) models in response to 800 ms current injections of varying stimulus amplitudes (indicated to the left of each panel). Parameters for the yPC: *a*_*NaT*_ = 1300, *a*_*CaL*_ = 25, *a*_*DK*_ = 6000.0, *a*_*SK*_ = 1600, *r*_*w*_ = 1.8, *r*_*c*_ = 5e^−3^, *k*_*c*_ = 6e^−6^. All parameters for the aPC the same except *a*_*CaL*_ = 50.

### 3.3 Modeling age-related changes in burst firing

#### 3.3.1 Bursting in response to stimulation

Some CA1 PCs fire bursts instead of trains of spikes [82], especially in certain developmental periods [83]. Burst firing can be generated in the model with several different parameter combinations. For the following simulations, we modified several of the current amplitudes and select kinetics, all within physiological limits (see Table 3). Under this parameter regime, model PCs are silent at rest but burst if stimulated (Fig. 3).

To explore the effects of aging on bursting, we fixed all parameters except for the maximum Ca^2+^ current amplitude, as previously. We then stimulated the two model PCs with a series of square-pulse current injections of increasing amplitudes (50-170 pA) to compare their responses. Again, the larger Ca^2+^ current in the aPC causes it to fire either sooner or nearly simultaneously with the yPC shortly after stimulus onset in all simulations. However, the relative timing of the PCs’ firing after the first burst depends on the stimulus amplitude (Fig. 3).

At lower stimulation amplitudes (50 and 80 pA; first two panels), the aPC continues to burst sooner or nearly simultaneously, but always fires fewer spikes per burst than the yPC (2 versus 3, respectively). As the stimulation amplitude increases (110 pA; 3rd panel), the two PCs again fire nearly simultaneously at the onset. However, because the aPC fires fewer spikes per burst, it is able to recover sooner and burst before the yPC for four cycles. It is only towards the end of the stimulus that the larger AHP in the aPC eventually brings it into sync again with the yPC. Finally, at the highest stimulation amplitudes (140 and 170 pA; 4th and 5th panels), both PCs show stronger adaptation, but the yPC maintains something akin to bursting, while the aPC ‘devolves’ to a pattern more like tonic spiking. Depending on what is considered the relevant electrical event – the single spike or the burst – the aPC shows an increased number of events relative to the yPC (13 spikes versus 8 burst-like events, respectively), but the normal bursting pattern is lost.

#### 3.3.2 Spontaneous bursting

A small percentage of CA1 PCs fire bursts in the absence of stimulation [84]. To generate this pattern, we again modifed several of the current amplitudes and select kinetics, all within physiological limits (see Table 3). Under this parameter regime, the yPC fires spontaneous bursts at a frequency of ∼1 Hz with 3 spikes per burst (Fig. 4, 3rd panel), similar to recordings [85].

**Figure 4:**
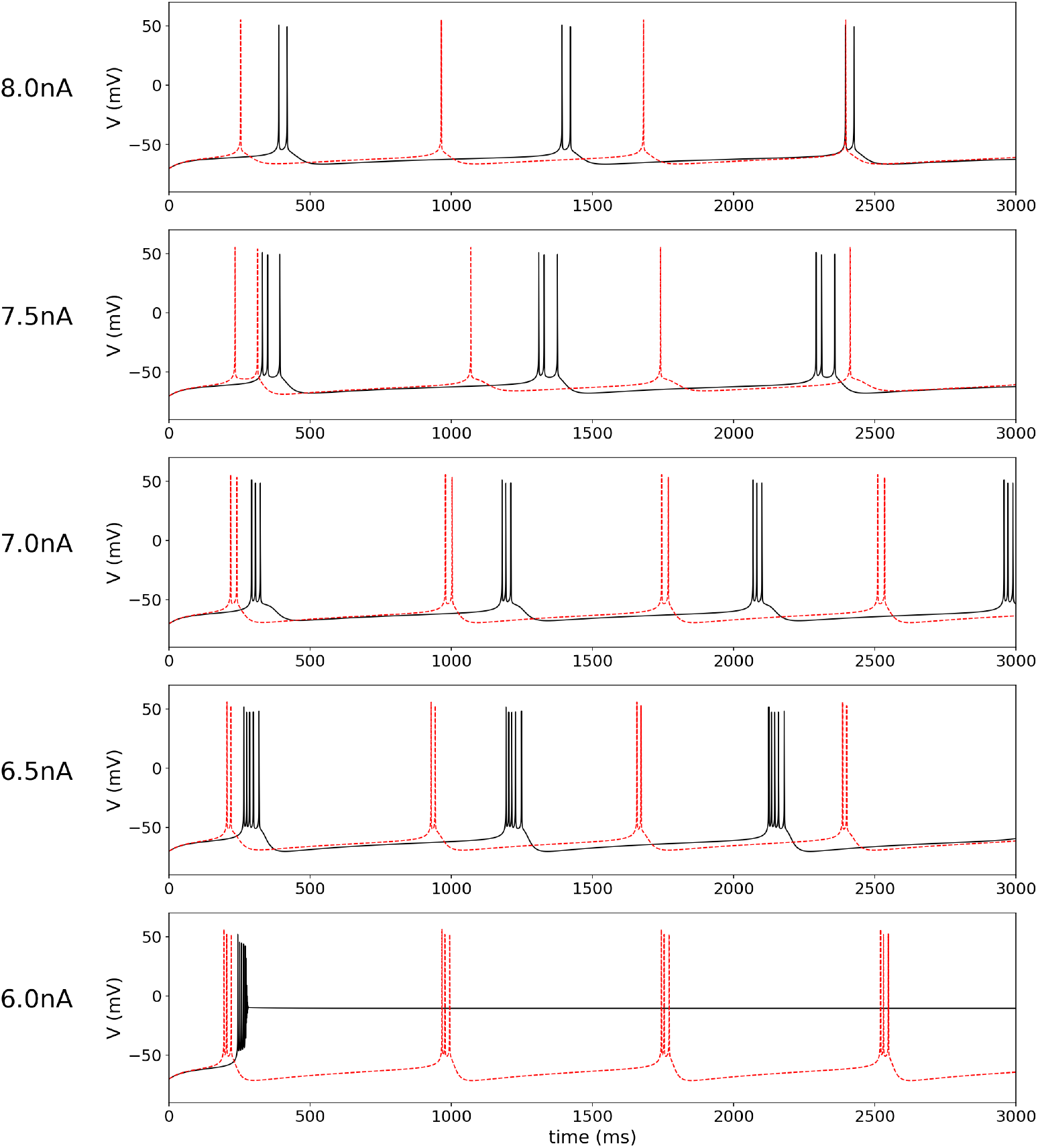
Comparison of spontaneous electrical activity in the yPC (solid black traces) and aPC (dashed red traces) for different levels of DK current amplitude. *a*_*DK*_ decreasing from 8000 to 6000 pA (8 to 6 nA) in steps of 500 pA, as indicated. Parameters: *a*_*NaT*_ = 2300, *a*_*CaL*_ = 25, *A*_*DK*_ = 7000, *a*_*SK*_ = 300, *r*_*w*_ = 1.1, *r*_*c*_ = 5e^−2^, *k*_*c*_ = 6e^−6^, *I*_*F*_ = 0.0. All parameters for aPC the same except *a*_*CaL*_ = 50.

Increasing the Ca^2+^ current, as previously, changed the spontaneous firing pattern (Fig. 4). Additional interesting effects can be seen if we vary the DK current amplitude within the range previously used for simulations, 6000-8000 pA (6-8 nA). At the highest DK amplitude, the yPC bursts spontaneously but with only 2 spikes per burst (Fig. 4, 1st panel). The aPC, however, does not burst but spikes tonically at a frequency of ∼2 Hz. When the DK amplitude is reduced (7.5 nA), the yPC continues to burst, now with 3 spikes per burst, while the aPC still spikes tonically.

Reducing the DK amplitude further (7 nA or 6.5 nA), causes both PCs to burst spontaneously, though the aPC always fires fewer spikes per burst. One can use our Jupyter notebook to explore further, and see that while the Ca^2+^ current is larger for the aPC, the maximum accumulation of intracellular Ca^2+^ is either similar or higher in the yPC due to its additional spiking. This causes a similar or larger SK current in the yPC, and results in a slower burst frequency.

Finally, reducing the DK amplitude to the lowest level (6.0 nA) removes more of the ‘brake’ on the yPC and causes it to spike at high frequency and then quickly block depolarize. The aPC, on the other hand, retains the spontaneous bursting pattern, now with more spikes per burst.

### 3.4 Responses to local field potential forcing

Square-pulse stimulation is useful for examining the timing of PC responses, but is not a physiologically realistic stimulus. Instead, to simulate local field potential (LFP) forcing onto CA1 PCs, we use an Ornstein-Uhlenbeck stochastic process [47, 48], as in Eq. 9. First, we reset the model with the parameters needed to produce adaptive firing, as in Section 3.1. In the yPC, LFP forcing produces repetitive, irregular firing at a frequency of ∼3 Hz on average (Fig. 5, top panel), which is similar to recordings of spontaneous firing in CA1 PCs [86], particularly in response to specific brain rhythms recorded in the surrounding field [87, 88].

**Figure 5:**
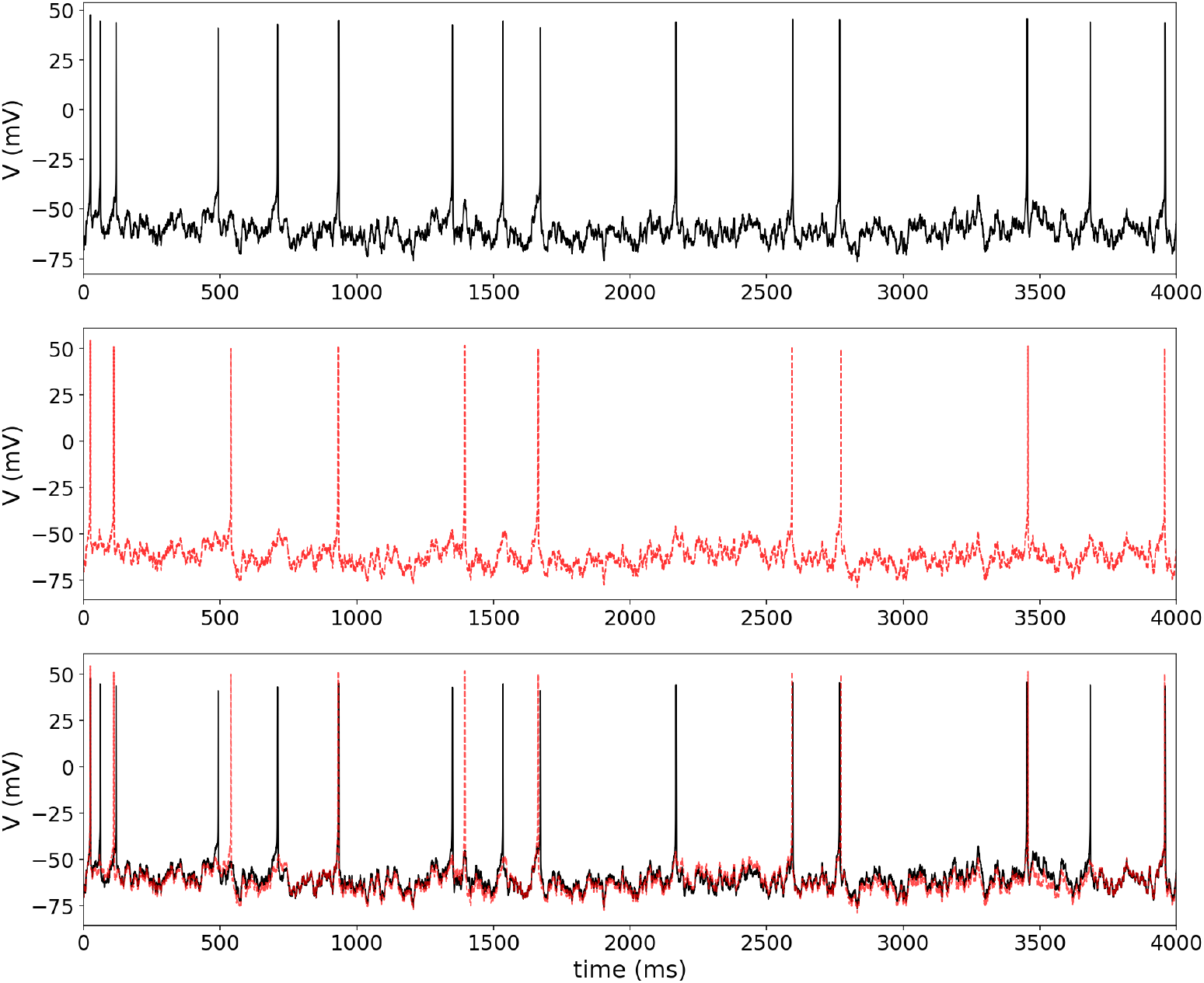
Responses of the yPC (solid black traces) and aPC (dashed red traces) to the same LFP forcing while in adaptive firing mode. Top panel shows just the yPC response, the 2nd panel shows the aPC response, and the 3rd panel shows the overlap of the two traces. Parameters for the yPC and aPC are the same as in Fig. 1. LFP parameters: *μ*_*F*_ =50.0 pA, *σ*_*F*_ =25.0 pA, *τ*_*F*_ =1/2.0 for both models.

In response to the exact same LFP forcing, the aPC with increased Ca^2+^ current shows a similar irregular firing pattern, but slower frequency of ∼2 Hz (Fig. 5, 2nd panel; compare overlap in 3rd panel). The simulation also shows several time points when the two cells fire almost simultaneously, and then the yPC fires again while the aPC does not. This apparent ‘spike failure’ has been seen in recordings of PCs from aged animals [13].

Next, we set the parameters to produce conditional bursting, as previously. Under these conditions, LFP forcing in the yPC produced irregular burst firing at a frequency of ∼5Hz (i.e. theta frequency) with 2-4 spikes per burst (Fig. 6, top panel). This firing pattern is similar to spontaneous activity recorded in a subset of CA1 PCs known as phasic theta-ON cells, which preferentially burst during theta activity recorded from the surrounding field [87, 89]. Increased Ca^2+^ current in the aPC changes the firing pattern (Fig. 6, 2nd panel). In response to LFP forcing, the aPC still fires irregular bursts, but with fewer spikes per burst (3 max) and a higher occurrence of 2-spike bursts than seen in the yPC. In addition, the aPC fires single APs amidst the bursts, which occurs half as frequently in the yPC under this parameter regime. Also, the timing of the bursts in the aPC is altered relative to the yPC (see 3rd panel overlap).

**Figure 6:**
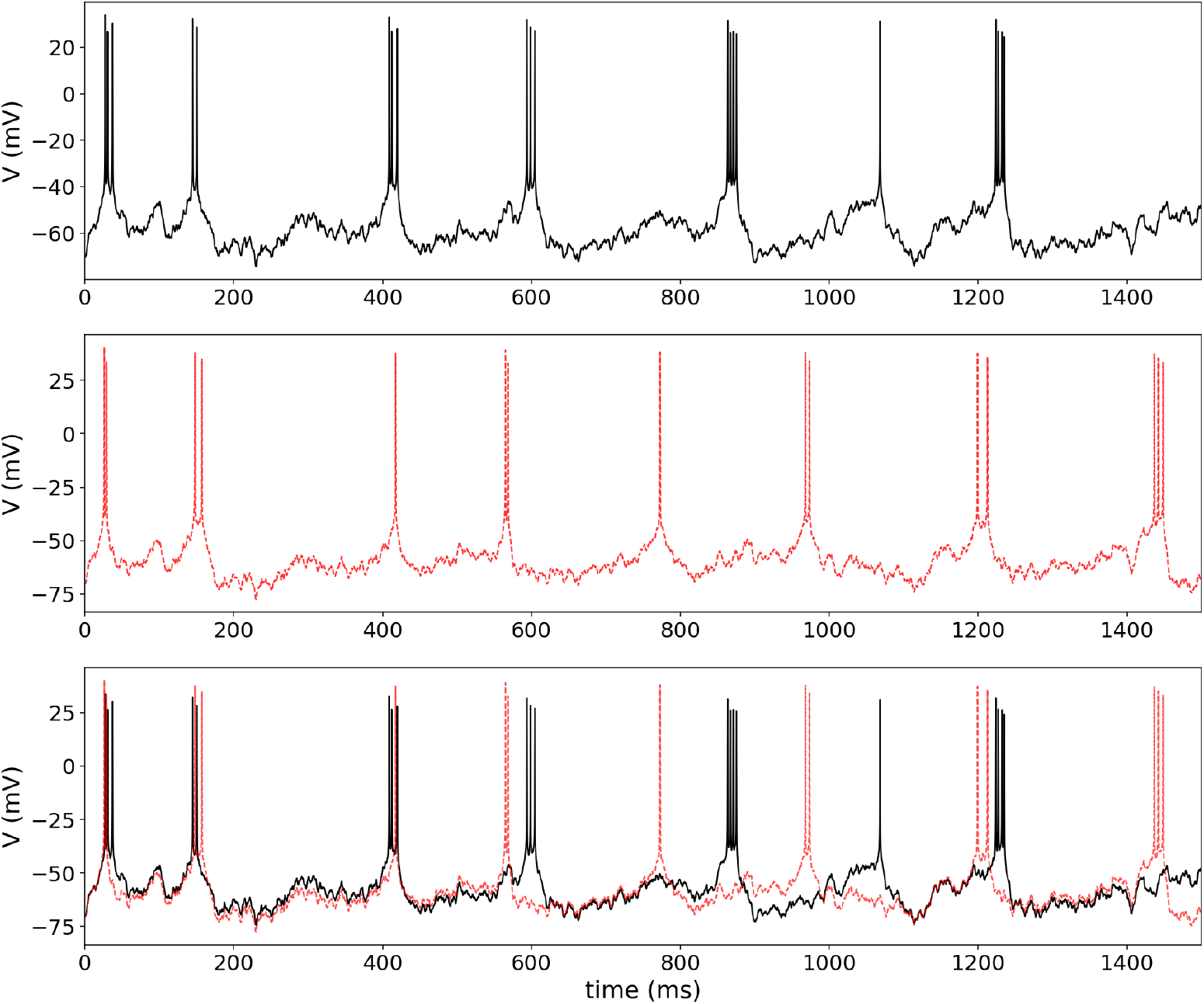
Responses of the conditionally bursting yPC (solid black traces) and aPC (dashed red traces) models to the same LFP forcing. Top panel shows just the yPC response, the 2nd panel shows the aPC response, and the 3rd panel shows the overlap of the two traces. Parameters for the yPC and aPC are the same as in Fig. 3. LFP parameters the same as in Fig. 5 except *σ*_*F*_ =20.0 pA.

## 4 Discussion

### 4.1 Cellular heterogeneity

Our three-dimensional, single-compartment model derived from first principles of thermodynamics can reproduce the diversity of firing patterns recorded in CA1 PCs. Moving between the different patterns was achieved primarily by changes to the relative expression of ion channels in the model. We did not systematically explore the full parameter space, but future work could include bifurcation analysis to determine boundaries for each firing pattern. The flexibility of the model could be useful for researchers to study the effects of PC heterogeneity on network function. Geiller and colleagues [90] write, “Until very recently, hippocampus models and theories were built on a view of homogenous population of principal cells” (pg. 6). However, there is heterogeneity in CA1 PCs, especially during different development stages (for review, see [31]). Lee and colleagues [91] write, “how the heterogeneous PCs integrate into the CA1 circuit remains unknown” (pg. 1129).

Experimentally, it is difficult to quantify how many PCs in a given network are displaying a specific firing pattern, and even harder, if not impossible, to manipulate these percentages. Furthermore, cells can transition between firing patterns [92], meaning percentages might fluctuate. With our minimal model, however, we could build small networks with different balances of adapting versus bursting PCs, and explore how this affects network output. We could also model the progression of aging in the network by varying the percentage of PCs which have altered Ca^2+^ channel density, or implement a whole spectrum of channel expression across the simulated network.

### 4.2 Aging and Ca^2+^ channel expression

Our model can also reproduce changes in electrical activity seen in aged CA1 PCs, including larger AHPs [12–14] and increased adaptation [17–19], by increasing the L-type Ca^2+^ current amplitude to a level similar to that found in recordings. The L-type channel was modeled after the Ca_*v*_1.2 isoform based on work showing this is the primary contributor in rodent brain, responsible for ∼70-80% of the L-type current [50, 93]. mRNA expression of *Cacnac1C* (the gene encoding Ca_*v*_1.2) is increased in aged mice and rats [22, 94]. Increases in plasma membrane expression [25] and phosphorylation [95] of Ca_*v*_1.2 channels have also been seen in aged rats. In addition, changes in *Cacna1c*/Ca_*v*_1.2 expression are correlated with memory impairments [94, 96].

However, CA1 PCs also express the Ca_*v*_1.3 isoform [97], which is responsible for ∼20% of the total L-type current [50, 93]. Studies have found both increased mRNA [22] and protein [24] expression of Ca_*v*_1.3 in aged rats, and this increased expression is correlated with memory impairment [98]. Knockout studies in mice indicate that it is this isoform, and not Ca_*v*_1.2, which contributes to slow AHP generation [99], possibly via activation of co-localized SK channels [97]. While experimental studies have been complicated by a lack of pharmacological agents which can isolate currents carried by the different isoforms, it would be relatively simple with our model to study the contributions of these two channels. The primary difference between the two is a shift in the activation curve of the Ca_*v*_1.3 channel to more hyperpolarized values, relative to Ca_*v*_1.2 [66]. Changing the parameter *v*_*j*_ in Eq. 4 would allow us to represent the different isoforms and explore how changes in the expression of each during aging might affect PC activity.

There are many cellular changes apart from Ca^2+^ channel expression that occur during aging and could contribute to altered activity in PCs. Studies have implicated Ca^2+^ release from intracellular stores as an important contributor, particularly to larger AHPs in aged animals (for reviews see [100, 101]). We did not explore the role of intracellular Ca^2+^ stores in this study, nor many of the other cellular changes that surely contribute to the multifactorial process of aging. It is not our intention to suggest Ca^2+^ channel expression is the only factor altering the activity of PCs in aged animals. Nevertheless, we do demonstrate that an increase in L-type Ca^2+^ channels is *sufficient* to reproduce many of the changes in PC firing seen during aging. These results agree with a previous modeling study, which also found that an increase in L-type Ca^2+^ conductance was sufficient to produce changes in adaptation and AHPs similar to those seen in aged CA1 PCs [102]. However, their model was mathematically complex, with 183 compartments and more than a dozen ionic currents – very different from the single-compartment, minimal model we present here. In addition, their study looked only at adaptation and AHPs, while our study goes further to investigate conditional bursting, spontaneous bursting, and responses to LFP stimulation.

### 4.3 Aging and excitability

Our simulations do show changes in the electrical activity of aged PCs, but do these changes represent decreased excitability? This question relates more broadly to how we think about excitability – the term is rarely clearly defined or used in a standardized way. In some studies, excitability is used to refer to a change in the firing rate of a cell over the course of an injected current pulse, claiming that PCs with stronger adaptation are less excitable [18]. In our simulations under the adaptive firing parameter regime, the aPC did have stronger adaptation and fired fewer times during the stimulation period than the yPC. However, under some circumstances, this stronger adaptation occurred only after the aPC initially fired faster than the yPC (see for example Fig. 2 inset). Should we consider this decreased excitability?

If what concerns us with excitability is the activity of the cell over a given time period, then the results under the bursting parameter regimes are even less clear. The aPC always fired fewer APs per burst than the yPC, indicating something akin to stronger adaptation. However, if the ‘event’ we are considering is instead the burst, there are conditions under which the aPC fired a greater number of bursts in a given time period than the yPC. How should we interpret these results with respect to excitability? To our knowledge, there are very few experimental studies to date that have compared burst firing in young versus aged animal CA1 PCs, perhaps because of the relatively low percentage of cells with this firing pattern in certain developmental periods [83]. The only study of which we are aware compared spontaneous burst firing in young versus aged animal PCs during rest and exploratory behavior [103]. Unfortunately, there are several factors that make it difficult to compare our results with theirs. First, their recordings were extracellular. As such, no bursts from individual PCs are shown in the paper to compare to our simulations, which reproduce intracellular firing. Furthermore, the study focused on interspike intervals (ISIs), rather than the overall bursting pattern (e.g. number of bursts in a given period), which was our focus. Interestingly, Smith and colleagues [103] found that while ISIs were more left-skewed in aged animals when burst firing was recorded at rest, they found no difference between the age groups during behavior, suggesting that compensatory mechanisms work differently during behavior to adjust for changes in cellular excitability. Such compensatory mechanisms could be explored using our model, for example by adding simulated cholinergic input.

Other researchers use the term excitability to describe how easy it is to get a cell to fire in response to stimulation, referring to “propensity” [104] or “readiness” [105]. In this context, excitability could be measured by the rheobase, or minimum current which generates firing in a neuron, as done in some studies of aging in CA1 [106]. However, ‘propensity’ or ‘readiness’ could also be interpreted as how quickly a cell fires after stimulus onset. In the AHP simulations, we saw that the aPC required additional current to fire the same number of spikes as for the yPC. On the other hand, the aPC often fired sooner than the yPC. These effects were a result of the increased Ca^2+^ current in aged cells – the larger Ca^2+^ current depolarized the cells faster and caused them to fire sooner, but it also caused the SK current to be larger and consequently slowed firing. It is as if the aPCs were initially more excitable, but then ‘burned out’ more quickly than the yPCs.

Overall, we believe our results highlight the importance of moving away from vague, ill-defined terms like ‘excitability’ in favor of precise language that describes the effect of interest (e.g., PCs spiked faster, spiked sooner, etc.). We hope that, in a broader sense, our study will encourage neuroscientists, and particularly aging researchers, to reevaluate how they think and write about excitability.

## References

[1] M.M. Oh, F.A. Oliveira, and J.F. Disterhoft. Learning and aging related changes in intrinsic neuronal excitability. Frontiers in Aging Neuroscience, 2:2, 2010. 10.3389/neuro.24.002.2010.

[2] E.S. Rosenzweig and C.A. Barnes. Impact of aging on hippocampal function: plasticity, network dynamics, and cognition. Progress in Neurobiology, 69(3):143–179, 2003. 10.1016/S0301-0082(02)00126-0.

[3] A.M. Fjell, L. McEvoy, D. Holland, A.M. Dale, K.B. Walhovd, and for the Alzheimer’s Disease Neuroimaging Initiative. What is normal in normal aging? Effects of aging, amyloid and Alzheimer’s disease on the cerebral cortex and the hippocampus. Progress in Neurobiology, 117:20–40, 2014. 10.1016/j.pneurobio.2014.02.004.

[4] M. Rodriguez, C. Rodriguez-Sabate, I. Morales, A. Sanchez, and M. Sabate. Parkinson’s disease as a result of aging. Aging Cell, 14(3):293–308, 2015. 10.1111/acel.12312.

[5] C.A. Barnes and B.L. McNaughton. An age comparison of the rates of acquisition and forgetting of spatial information in relation to long-term enhancement of hippocampal synapses. Behavioral Neuroscience, 99(6):1040, 1985. 10.1037/0735-7044.99.6.1040.

[6] F.H. Gage, S.B. Dunnett, and A. Björklund. Spatial learning and motor deficits in aged rats. Neurobiology of Aging, 5(1):43–48, 1984. 10.1016/0197-4580(84)90084-8.

[7] A. Caprioli, O. Ghirardi, A. Giuliani, M.T. Ramacci, and L. Angelucci. Spatial learning and memory in the radial maze: A longitudinal study in rats from 4 to 25 months of age. Neurobiology of Aging, 12(5):605–607, 1991. 10.1016/0197-4580(91)90093-Y.

[8] S.M. Wilkniss, M.G. Jones, D.L. Korol, P.E. Gold, and C.A. Manning. Age-related differences in an ecologically based study of route learning. Psychology and Aging, 12(2):372, 1997. 10.1037/0882-7974.12.2.372.

[9] D.L. Deupree, J. Bradley, and D.A. Turner. Age-related alterations in potentiation in the CA1 region in F344 rats. Neurobiology of Aging, 14(3):249–258, 1993. 10.1016/0197-4580(93)90009-Z.

[10] E.S. Rosenzweig, G. Rao, B.L. McNaughton, and C.A. Barnes. Role of temporal summation in age-related long-term potentiation–induction deficits. Hippocampus, 7(5):549–558, 1997. 10.1002/(SICI)1098-1063(1997)7:5<549::AID-HIPO10>3.0.CO;2-0.

[11] P.W. Landfield, J.L. McGaugh, and G. Lynch. Impaired synaptic potentiation processes in the hippocampus of aged, memory-deficient rats. Brain Research, 150(1):85–101, 1978. 10.1016/0006-8993(78)90655-8.

[12] P.W. Landfield and T.A. Pitler. Prolonged Ca^2+^-dependent afterhyperpolarizations in hippocampal neurons of aged rats. Science, 226(4678):1089–1092, 1984. 10.1126/science.6494926.

[13] J.C. Gant and O. Thibault. Action potential throughput in aged rat hippocampal neurons: regulation by selective forms of hyperpolarization. Neurobiology of Aging, 30(12):2053–2064, 2009. 10.1016/j.neurobiolaging.2008.02.006.

[14] J.M. Power, W.W. Wu, E. Sametsky, M.M. Oh, and J.F. Disterhoft. Age-related enhancement of the slow outward calcium-activated potassium current in hippocampal CA1 pyramidal neurons in vitro. Journal of Neuroscience, 22(16):7234–7243, 2002. 10.1523/JNEUROSCI.22-16-07234.2002.

[15] B.E. Alger and R.A. Nicoll. Epileptiform burst afterhyperolarization: Calcium-dependent potassium potential in hippocampal CA1 pyramidal cells. Science, 210(4474):1122–1124, 1980. 10.1126/science.7444438.

[16] J.R. Hotson and D.A. Prince. A calcium-activated hyperpolarization follows repetitive firing in hippocampal neurons. Journal of Neurophysiology, 43(2):409–419, 1980. 10.1152/jn.1980.43.2.409.

[17] J.C. Gant, M.M. Sama, P.W. Landfield, and O. Thibault. Early and simultaneous emergence of multiple hippocampal biomarkers of aging is mediated by Ca2+-induced Ca2+ release. Journal of Neuroscience, 26(13):3482–3490, 2006. 10.1523/JNEUROSCI.4171-05.2006.

[18] J.R. Moyer, L.T. Thompson, J.P. Black, and J.F. Disterhoft. Nimodipine increases excitability of rabbit CA1 pyramidal neurons in an age- and concentration-dependent manner. Journal of Neurophysiology, 68(6):2100–2109, 1992. 10.1152/jn.1992.68.6.2100.

[19] G.C. Tombaugh, W.B. Rowe, and G.M. Rose. The slow afterhyperpolarization in hippocampal CA1 neurons covaries with spatial learning ability in aged Fisher 344 rats. Journal of Neuroscience, 25(10):2609–2616, 2005. 10.1523/JNEUROSCI.5023-04.2005.

[20] L.W. Campbell, S-Y. Hao, O. Thibault, E.M. Blalock, and P.W. Landfield. Aging changes in voltagegated calcium currents in hippocampal CA1 neurons. Journal of Neuroscience, 16(19):6286–6295, 1996. 10.1523/JNEUROSCI.16-19-06286.1996.

[21] O. Thibault, R. Hadley, and P.W. Landfield. Elevated postsynaptic [Ca2+]i and L-type calcium channel activity in aged hippocampal neurons: Relationship to impaired synaptic plasticity. Journal of Neuroscience, 21(24):9744–9756, 2001. 10.1523/JNEUROSCI.21-24-09744.2001.

[22] J.P. Herman, K-C. Chen, R. Booze, and P.W. Landfield. Up-regulation of α1D Ca2+ channel subunit mRNA expression in the hippocampus of aged F344 rats. Neurobiology of Aging, 19(6):581–587, 1998. 10.1016/S0197-4580(98)00099-2.

[23] O. Thibault and P.W. Landfield. Increase in single L-type calcium channels in hippocampal neurons during aging. Science, 272(5264):1017–1020, 1996. 10.1126/science.272.5264.1017.

[24] L.M. Veng and M.D. Browning. Regionally selective alterations in expression of the α1D subunit (Cav 1.3) of L-type calcium channels in the hippocampus of aged rats. Molecular Brain Research, 107(2):120–127, 2002. 10.1016/S0169-328X(02)00453-9.

[25] F.L. Núñez-Santana, M.M. Oh, M.D. Antion, A. Lee, J.W. Hell, and J.F. Disterhoft. Surface L-type Ca^2+^ channel expression levels are increased in aged hippocampus. Aging Cell, 13(1):111–120, 2014. 10.1111/acel.12157.

[26] C.M. Norris, S. Halpain, and T.C. Foster. Reversal of age-related alterations in synaptic plasticity by blockade of L-type Ca2+ channels. Journal of Neuroscience, 18(9):3171–3179, 1998. 10.1523/JNEUROSCI.18-09-03171.1998.

[27] M. Sandin, S. Jasmin, and T.E. Levere. Aging and cognition: Facilitation of recent memory in aged nonhuman primates by nimodipine. Neurobiology of Aging, 11(5):573–575, 1990. 10.1016/0197-4580(90)90120-O.

[28] M.A. Herrera-Valdez. Membranes with the same ion channel populations but different excitabilities. PLOS ONE, 7(4):e34636, 2012. 10.1371/journal.pone.0034636.

[29] M.A. Herrera-Valdez, E.C. McKiernan, S.D. Berger, S. Ryglewski, C. Duch, and S. Crook. Relating ion channel expression, bifurcation structure, and diverse firing patterns in a model of an identified motor neuron. Journal of Computational Neuroscience, 34(2):211–229, 2013. 10.1007/s10827-012-0416-6.

[30] M.A. Herrera-Valdez. A thermodynamic description for physiological transmembrane transport [version 2; peer review: 2 approved]. F1000Research, 7, 2018. 10.12688/f1000research.16169.2.

[31] E.C. McKiernan and D.F. Marrone. CA1 pyramidal cells have diverse biophysical properties, affected by development, experience, and aging. PeerJ, 5:e3836, 2017. 10.7717/peerj.3836.

[32] J.L. Hindmarsh and R.M. Rose. A model of neuronal bursting using three coupled first order differential equations. Proceedings of the Royal Society B, 221(1222):87–102, 1984. 10.1098/rspb.1984.0024.

[33] E. Av-Ron, H. Parnas, and L.A. Segel. A basic biophysical model for bursting neurons. Biological Cybernetics, 69(1):87–95, 1993. 10.1007/BF00201411.

[34] J. Rinzel. Excitation dynamics: insights from simplified membrane models. Federation Proceedings, 44(15):2944–2946, 1985. PMID: 2415401.

[35] E. Av-Ron, H. Parnas, and L. A. Segel. A minimal biophysical model for an excitable and oscillatory neuron. Biological Cybernetics, 65(6):487–500, 1991. 10.1007/BF00204662.

[36] M.A. Herrera-Valdez. An equation for the biological transmembrane potential from basic biophysical principles [version 1; peer review: 2 not approved]. F1000Research, 9:676, 2020. 10.12688/f1000research.24205.1.

[37] L. Groc, Z. Petanjek, B. Gustafsson, Y. Ben-Ari, E. Hanse, and R. Khazipov. In vivo blockade of neural activity alters dendritic development of neonatal CA1 pyramidal cells. European Journal of Neuroscience, 16(10):1931–1938, 2002. 10.1046/j.1460-9568.2002.02264.x.

[38] B. Hille. Ion Channels of Excitable Membranes. Sinauer Associates, Sunderland, Massachusetts, U.S.A, 3rd edition, 2001.

[39] M. Stocker. Ca2+-activated K+ channels: Molecular determinants and function of the SK family. Nature Reviews Neuroscience, 5:758–770, 2004. 10.1038/nrn1516.

[40] M.I. Kalinin and S.A. Kononogov. Boltzmann’s constant, the energy meaning of temperature, and thermodynamic irreversibility. Measurement Techniques, 48(7):632–636, 2005. 10.1007/s11018-005-0195-9.

[41] M. Covarrubias, A. Wei, and L. Salkoff. Shaker, Shal, Shab, and Shaw expresss independent K^+^-current systems. Neuron, 7(5):763–773, 1991. 10.1016/0896-6273(91)90279-9.

[42] A.L. Hodgkin and A.F. Huxley. A quantitative description of membrane current and its application to conduction and excitation in nerve. The Journal of Physiology, 117(4):500–544, 1952. 10.1113/jphysiol.1952.sp004764.

[43] S. Tsunoda and L. Salkoff. The major delayed rectifier in both Drosophila neurons and muscle is encoded by Shab. Journal of Neuroscience, 15(7):5209–5221, 1995. 10.1523/JNEUROSCI.15-07-05209.1995.

[44] B. Hirschberg, J. Maylie, J.P. Adelman, and N.V. Marrion. Gating of recombinant small-conductance Ca-activated K+ channels by calcium. Journal of General Physiology, 111(4):565–581, 1998. 10.1085/jgp.111.4.565.

[45] M. Rudolph and A. Destexhe. Characterization of subthreshold voltage fluctuations in neuronal membranes. Neural Computation, 15(11):2577–2618, 2003. 10.1162/089976603322385081.

[46] D.T. Gillespie. Exact numerical simulation of the Ornstein-Uhlenbeck process and its integral. Physical Review E, 54(2):2084, 1996. 10.1103/PhysRevE.54.2084.

[47] M. Rudolph, Z. Piwkowska, M. Badoual, T. Bal, and A. Destexhe. A method to estimate synaptic conductances from membrane potential fluctuations. Journal of Neurophysiology, 91(6):2884–2896, 2004. 10.1152/jn.01223.2003.

[48] A. Destexhe, M. Badoual, Z. Piwkowska, T. Bal, and M. Rudolph. A novel method for characterizing synaptic noise in cortical neurons. Neurocomputing, 58:191–196, 2004. 10.1016/j.neucom.2004.01.042.

[49] H. Murakoshi and J.S. Trimmer. Identification of the Kv2.1 K+ channel as a major component of the delayed rectifier K+ current in rat hippocampal neurons. Journal of Neuroscience, 19(5):1728–1735, 1999. 10.1523/JNEUROSCI.19-05-01728.1999.

[50] J.W. Hell, R.E. Westenbroek, C. Warner, M.K. Ahlijanian, W. Prystay, M.M. Gilbert, T.P. Snutch, and W.A. Catterall. Identification and differential subcellular localization of the neuronal class C and class D L-type calcium channel alpha 1 subunits. Journal of Cell Biology, 123(4):949–962, 1993. 10.1083/jcb.123.4.949.

[51] M. Li, J.W. West, Y. Lai, T. Scheuer, and W.A. Catterall. Functional modulation of brain sodium channels by cAMP-dependent phosphorylation. Neuron, 8(6):1151–1159, 1992. 10.1016/0896-6273(92)90135-Z.

[52] D.A. Fadool and I.B. Levitan. Modulation of olfactory bulb neuron potassium current by tyrosine phosphorylation. Journal of Neuroscience, 18(16):6126–6137, 1998. 10.1523/JNEUROSCI.18-16-06126.1998.

[53] S.O.M. Ketelaars, J.A. Gorter, E.A. van Vliet, F.H. Lopes da Silva, and W.J. Wadman. Sodium currents in isolated rat CA1 pyramidal and dentate granule neurones in the post-status epilepticus model of epilepsy. Neuroscience, 105(1):109–120, 2001. 10.1016/S0306-4522(01)00176-2.

[54] P. Sah, A.J. Gibb, and P.W. Gage. The sodium current underlying action potentials in guinea pig hippocampal CA1 neurons. Journal of General Physiology, 91(3):373–398, 1988. 10.1085/jgp.91.3.373.

[55] D.P. Mohapatra, H. Misonou, P. Sheng-Jun, J.E. Held, D.J. Surmeier, and J.S. Trimmer. Regulation of intrinsic excitability in hippocampal neurons by activity-dependent modulation of the KV 2.1 potassium channel. Channels, 3(1):46–56, 2009. 10.4161/chan.3.1.7655.

[56] J. Scuvée-Moreau, A. Boland, A. Graulich, L. Van Overmeire, D. D’hoedt, F. Graulich-Lorge, E. Thomas, A. Abras, M. Stocker, J-F. Liégeois, and V. Seutin. Electrophysiological characterization of the SK channel blockers methyl-laudanosine and methyl-noscapine in cell lines and rat brain slices. British Journal of Pharmacology, 143(6):753–764, 2004. 10.1038/sj.bjp.0705979.

[57] K.S. Richards, K. Bommert, G. Szabo, and R. Miles. Differential expression of Na+/K+-ATPase α-subunits in mouse hippocampal interneurones and pyramidal cells. The Journal of Physiology, 585(2):491–505, 2007. 10.1113/jphysiol.2007.144733.

[58] D. Johnston and S. M-S. Wu. Foundations of Cellular Neurophysiology. MIT Press, 1995.

[59] D.M. Egelman and P.R. Montague. Calcium dynamics in the extracellular space of mammalian neural tissue. Biophysical Journal, 76(4):1856–1867, 1999. 10.1016/S0006-3495(99)77345-5.

[60] L.P. Endresen, K. Hall, J.S. Høye, and J. Myrheim. A theory for the membrane potential of living cells. European Biophysics Journal, 29(2):90–103, 2000. 10.1007/s002490050254.

[61] P.D. Weer, D.C. Gadsby, and R.F. Rakowski. Voltage dependence of the Na-K pump. Annual Review of Physiology, 50(1):225–241, 1988. 10.1146/annurev.ph.50.030188.001301.

[62] B.C. Carter and B.P. Bean. Sodium entry during action potentials of mammalian neurons: incomplete inactivation and reduced metabolic efficiency in fast-spiking neurons. Neuron, 64(6):898–909, 2009. 10.1016/j.neuron.2009.12.011.

[63] S. Gasparini and J.C. Magee. Phosphorylation-dependent differences in the activation properties of distal and proximal dendritic Na+ channels in rat CA1 hippocampal neurons. The Journal of Physiology, 541(3):665–672, 2002. 10.1113/jphysiol.2002.020503.

[64] M. Martina and P. Jonas. Functional differences in Na+ channel gating between fast-spiking interneurones and principal neurones of rat hippocampus. The Journal of Physiology, 505(3):593–603, 1997. 10.1111/j.1469-7793.1997.593ba.x.

[65] J.C. Magee and D. Johnston. Characterization of single voltage-gated Na+ and Ca2+ channels in apical dendrites of rat CA1 pyramidal neurons. The Journal of Physiology, 487(1):67, 1995. 10.1113/jphysiol.1995.sp020862.

[66] W. Xu and D. Lipscombe. Neuronal Cav 1.3 α1 L-type channels activate at relatively hyperpolarized membrane potentials and are incompletely inhibited by dihydropyridines. Journal of Neuroscience, 21(16):5944–5951, 2001. 10.1523/JNEUROSCI.21-16-05944.2001.

[67] B. Balasubramanian, J.P. Imredy, D. Kim, J. Penniman, A. Lagrutta, and J.J. Salata. Optimization of Cav 1.2 screening with an automated planar patch clamp platform. Journal of Pharmacological and Toxicological Methods, 59(2):62–72, 2009. 10.1016/j.vascn.2009.02.002.

[68] B.C. Carter, A.J. Giessel, B.L. Sabatini, and B.P. Bean. Transient sodium current at subthreshold voltages: activation by epsp waveforms. Neuron, 75(6):1081–1093, 2012. 10.1016/j.neuron.2012.08.033.

[69] P.F. Costa. The kinetic parameters of sodium currents in maturing acutely isolated rat hippocampal CA1 neurones. Developmental Brain Research, 91(1):29–40, 1996. 10.1016/0165-3806(95)00159-X.

[70] S.K. Shin, H.Y. Li, K. Cho, Y.W. Cho, J-H. Lee, and K-S. Park. Sites and regulation of L-type Ca2+ channel Cav 1.2 phosphorylation in brain. Cellular and Molecular Neurobiology, pages 1–5, 2021. 10.1007/s10571-021-01095-2.

[71] A.M.J. VanDongen, G.C. Frech, J.A. Drewe, R.H. Joho, and A.M. Brown. Alteration and restoration of K+ channel function by deletions at the N-and C-termini. Neuron, 5(4):433–443, 1990. 10.1016/0896-6273(90)90082-Q.

[72] M.M. Oh, F.A. Oliveira, J. Waters, and J.F. Disterhoft. Altered calcium metabolism in aging CA1 hippocampal pyramidal neurons. Journal of Neuroscience, 33(18):7905–7911, 2013. 10.1523/JNEUROSCI.5457-12.2013.

[73] J.C. Magee, R.B. Avery, B.R. Christie, and D. Johnston. Dihydropyridine-sensitive, voltage-gated Ca2+ channels contribute to the resting intracellular Ca2+ concentration of hippocampal CA1 pyramidal neurons. Journal of Neurophysiology, 76(5):3460–3470, 1996. 10.1152/jn.1996.76.5.3460.

[74] C.R. Harris, K.J. Millman, S.J. van der Walt, R. Gommers, P. Virtanen, D. Cournapeau, E. Wieser, J. Taylor, S. Berg, N.J. Smith, R. Kern, M. Picus, S. Hoyer, M.H. van Kerkwijk, M. Brett, A. Haldane, J. Fernández del Río, M. Wiebe, P. Peterson, P. Gérard-Marchant, K. Sheppard, T. Reddy, W. Weckesser, H. Abbasi, C. Gohlke, and T.E. Oliphant. Array programming with NumPy. Nature, 585(7825):357–362, 2020. 10.1038/s41586-020-2649-2.

[75] J.D. Hunter. Matplotlib: A 2D graphics environment. Computing in Science & Engineering, 9(3):90–95, 2007. 10.1109/MCSE.2007.55.

[76] D. Mondaca. pyprocess, 2012–. https://github.com/Zincr0/pyprocess.

[77] T. Kluyver, B. Ragan-Kelley, F. Pérez, B.E. Granger, M. Bussonnier, J. Frederic, K. Kelley, J.B. Hamrick, J. Grout, S. Corlay, P. Ivanov, D. Avila, S. Abdalla, C. Willing, and Jupyter Development Team. Jupyter Notebooks-a publishing format for reproducible computational workflows. In F. Loizides and B. Schmidt, editors, Positioning and Power in Academic Publishing: Players, Agents and Agendas, pages 87–90, 2016. 10.3233/978-1-61499-649-1-87.

[78] N. Gu, H. Hu, K. Vervaeke, and J.F. Storm. SK (KCa2) channels do not control somatic excitability in CA1 pyramidal neurons but can be activated by dendritic excitatory synapses and regulate their impact. Journal of Neurophysiology, 100(5):2589–2604, 2008. 10.1152/jn.90433.2008.

[79] J.F. Disterhoft, L.T. Thompson, J.R. Moyer, and D.J. Mogul. Calcium-dependent afterhyperpolarization and learning in young and aging hippocampus. Life Sciences, 59(5):413–420, 1996. 10.1016/0024-3205(96)00320-7.

[80] J.F. Storm. An after-hyperpolarization of medium duration in rat hippocampal pyramidal cells. Journal of Physiology, 409(1):171–190, 1989. 10.1113/jphysiol.1989.sp017491.

[81] E.A. Matthews, J.M. Linardakis, and J.F. Disterhoft. The fast and slow afterhyperpolarizations are differentially modulated in hippocampal neurons by aging and learning. Journal of Neuroscience, 29(15):4750–4755, 2009. 10.1523/JNEUROSCI.0384-09.2009.

[82] H. Su, G. Alroy, E.D. Kirson, and Y. Yaari. Extracellular calcium modulates persistent sodium current-dependent burst-firing in hippocampal pyramidal neurons. Journal of Neuroscience, 21(12):4173–4182, 2001. 10.1523/JNEUROSCI.21-12-04173.2001.

[83] S. Chen, C. Yue, and Y. Yaari. A transitional period of Ca2+-dependent spike afterdepolarization and bursting in developing rat CA1 pyramidal cells. Journal of Physiology, 567(1):79–93, 2005. 10.1113/jphysiol.2005.084590.

[84] M.S. Jensen, R. Azouz, and Y. Yaari. Variant firing patterns in rat hippocampal pyramidal cells modulated by extracellular potassium. Journal of Neurophysiology, 71(3):831–839, 1994. 10.1152/jn.1994.71.3.831.

[85] D. Golomb, C. Yue, and Y. Yaari. Contribution of persistent Na+ current and M-type K+ current to somatic bursting in CA1 pyramidal cells: combined experimental and modeling study. Journal of Neurophysiology, 96(4):1912–1926, 2006. 10.1152/jn.00205.2006.

[86] F. Manseau and S. Williams. Tuning in the hippocampal theta band in vitro: methodologies for recording from the isolated rodent septohippocampal circuit. Journal of Visualized Experiments, 126:e55851, 2017. 10.3791/55851.

[87] B.H. Bland, J. Konopacki, and R. Dyck. Heterogeneity among hippocampal pyramidal neurons revealed by their relation to theta-band oscillation and synchrony. Experimental Neurology, 195(2):458–474, 2005. 10.1016/j.expneurol.2005.06.007.

[88] C.Y.L. Huh, B. Amilhon, K.A. Ferguson, F. Manseau, S.G. Torres-Platas, J.P. Peach, S. Scodras, N. Mechawar, F.K. Skinner, and S. Williams. Excitatory inputs determine phase-locking strength and spike-timing of CA1 stratum oriens/alveus parvalbumin and somatostatin interneurons during intrinsically generated hippocampal theta rhythm. Journal of Neuroscience, 36(25):6605–6622, 2016. 10.1523/JNEUROSCI.3951-13.2016.

[89] L.V. Colom and B.H. Bland. State-dependent spike train dynamics of hippocampal formation neurons: evidence for theta-on and theta-off cells. Brain Research, 422(2):277–286, 1987. 10.1016/0006-8993(87)90934-6.

[90] T. Geiller, S. Royer, and J-S. Choi. Segregated cell populations enable distinct parallel encoding within the radial axis of the CA1 pyramidal layer. Experimental Neurobiology, 26(1):1–10, 2017. 10.5607/en.2017.26.1.1.

[91] S-H. Lee, I. Marchionni, M. Bezaire, C. Varga, N. Danielson, M. Lovett-Barron, A. Losonczy, and Soltesz. Parvalbumin-positive basket cells differentiate among hippocampal pyramidal cells. Neuron, 82(5):1129–1144, 2014. 10.1016/j.neuron.2014.03.034.

[92] M. Steriade, I. Timofeev, N. Dürmüller, and F. Grenier. Dynamic properties of corticothalamic neurons and local cortical interneurons generating fast rhythmic (30–40 Hz) spike bursts. Journal of Neurophysiology, 79(1):483–490, 1998. 10.1152/jn.1998.79.1.483.

[93] M.J. Sinnegger-Brauns, A. Hetzenauer, I.G. Huber, E. Renström, G. Wietzorrek, S. Berjukov, M. Cavalli, D. Walter, A. Koschak, R. Waldschütz, S. Herin, S. Bova, P. Rorsman, O. Pongs, N. Singewald, and J. Striessnig. Isoform-specific regulation of mood behavior and pancreatic β cell and cardiovascular function by L-type Ca2+ channels. The Journal of Clinical Investigation, 113(10):1430–1439, 2004. 10.1172/JCI20208.

[94] P. Zanos, S. Bhat, C.E. Terrillion, R.J. Smith, L.H. Tonelli, and T.D. Gould. Sex-dependent modulation of age-related cognitive decline by the L-type calcium channel gene Cacna1c (Cav 1.2). European Journal of Neuroscience, 42(8):2499–2507, 2015. 10.1111/ejn.12952.

[95] M.A. Davare and J.W. Hell. Increased phosphorylation of the neuronal L-type Ca2+ channel Cav 1.2 during aging. Proceedings of the National Academy of Sciences, 100(26):16018–16023, 2003. 10.1073/pnas.2236970100.

[96] S. Moosmang, N. Haider, N. Klugbauer, H. Adelsberger, N. Langwieser, J. Müller, M. Stiess, E. Marais, V. Schulla, L. Lacinova, S. Goebbels, K.A. Nave, D.R. Storm, F. Hofmann, and T. Kleppisch. Role of hippocampal Cav 1.2 Ca2+ channels in NMDA receptor-independent synaptic plasticity and spatial memory. Journal of Neuroscience, 25(43):9883–9892, 2005. 10.1523/JNEUROSCI.1531-05.2005.

[97] S.E.H. Bowden, S. Fletcher, D.J. Loane, and N.V. Marrion. Somatic colocalization of rat SK1 and D class (Cav 1.2) L-type calcium channels in rat CA1 hippocampal pyramidal neurons. Journal of Neuroscience, 21(20):RC175–RC175, 2001. 10.1523/JNEUROSCI.21-20-j0006.2001.

[98] L.M. Veng, M.H. Mesches, and M.D. Browning. Age-related working memory impairment is correlated with increases in the L-type calcium channel protein α1D (Cav 1.3) in area CA1 of the hippocampus and both are ameliorated by chronic nimodipine treatment. Molecular Brain Research, 110(2):193–202, 2003. 10.1016/S0169-328X(02)00643-5.

[99] A.E. Gamelli, B.C. McKinney, J.A. White, and G.G. Murphy. Deletion of the L-type calcium channel Cav 1.3 but not Cav 1.2 results in a diminished sAHP in mouse CA1 pyramidal neurons. Hippocampus, 21(2):133–141, 2011. 10.1002/hipo.20728.

[100] O. Thibault, J.C. Gant, and P.W. Landfield. Expansion of the calcium hypothesis of brain aging and Alzheimer’s disease: minding the store. Aging Cell, 6(3):307–317, 2007. 10.1111/j.1474-9726.2007.00295.x.

[101] E.C. Toescu and M. Vreugdenhil. Calcium and normal brain ageing. Cell Calcium, 47(2):158–164, 2010. 10.1016/j.ceca.2009.11.013.

[102] M. Markaki, S. Orphanoudakis, and P. Poirazi. Modelling reduced excitability in aged CA1 neurons as a calcium-dependent process. Neurocomputing, 65:305–314, 2005. 10.1016/j.neucom.2004.10.023.

[103] A.C. Smith, J.L. Gerrard, C.A. Barnes, and B.L. McNaughton. Effect of age on burst firing characteristics of rat hippocampal pyramidal cells. NeuroReport, 11(17):3865–3871, 2000. 10.1097/00001756-200011270-00052.

[104] G. Daoudal and D. Debanne. Long-term plasticity of intrinsic excitability: learning rules and mechanisms. Learning & Memory, 10(6):456–465, 2003. 10.1101/lm.64103.

[105] L.M. Konstantinović and S.R. Filipović. Effects of near-infrared low-level laser stimulation on neuronal excitability. In M.R. Hamblin and Y-Y. Huang, editors, Photobiomodulation in the Brain: Low-Level Laser (Light) Therapy in Neurology and Neuroscience, pages 233–240. Academic Press, 2019. 10.1016/B978-0-12-815305-5.00018-X.

[106] B. Potier, Y. Lamour, and P. Dutar. Age-related alterations in the properties of hippocampal pyramidal neurons among rat strains. Neurobiology of Aging, 14(1):17–25, 1993. 10.1016/0197-4580(93)90016-5.

